# *Insilico* and *Invitro* optimization of Naringin and rutin molecules targeting DNA damage in breast cancer cells

**DOI:** 10.1101/2021.12.21.473577

**Authors:** Badhe Pravin, Vivek Nanaware, Badhe Ashwini

## Abstract

Discovering the molecular mechanisms of DNA damage response pathways has led to new therapeutic approaches in oncology. Our study optimized DNA damage-targeting molecules naringin and rutin in breast cancer cells.

Our study involved MTT assays for detection of its toxicity and proliferative activity in breast cancer cells and normal cancer cells. Our studies determined the molecules’ antioxidant properties using the DPPH assay. The role in reducing free radicals has been evaluated using a variety of free radical scavenging activity assays.

Further evaluation of the molecules was carried out by high alkaline comet assay (pH >13) to test for genotoxicity. Human Dermal Fibroblast cells (2DD) (1×10^5^ cells/ml) and breast cancer cells (MDA-MB-231) were pre-incubated with Naringin and Rutin (10 µM) for one hour.

In normal cells, rutin and naringin molecules do not cause genotoxicity, but they cause DNA damage in breast cancer cells when they are diluted to 10µM. The results from our study indicate that both molecules cause 60-70% DNA damage in breast cancer cells.

## Introduction

Cellular DNA damage occurs when physical or chemical changes occur in DNA, affecting the transmission and interpretation of genetic code. Damage to DNA can be caused by a variety of endogenous and exogenous insults, including chemicals, radiation, free radicals, and topological changes, resulting in distinct types of damage [1]. Exogenous damage to DNA is caused by X-rays, cosmic rays, UV radiation, and secondary pollutants caused by chemical oxidation [2, 3]. Endogenous DNA damage occurs when cellular signaling and metabolic pathways that are necessary for healthy living are disrupted [4, 5]. Studies suggest chronic oxidative stress conditions are strongly associated with cancer [6].

A DNA lesion occurs to the cells in the human body every day. If these lesions are not repaired or if they are repaired incorrectly, they can lead to mutations or wider-scale genome aberrations that threaten the viability of a cell or organism [7]. A continuous exposure of DNA and genome to damaging factors leads to a variety of genetic defects that might be inherited from one generation to the next [8]. A surplus of free radicals causes damage to important biomolecules in living organisms [9]. Free radicals and reactive oxygen species are formed by a number of factors. Change in life style, restless living, types of dietary elements, junk foods, and smoking are the major contributing factors [10].

Human cells are constantly exposed to endogenous and exogenous agents that can generate genomic instability. These can lead to structural damage to DNA that can alter or eliminate fundamental cellular processes, such as DNA replication or transcription. They can also produce base and sugar modifications, induce single and double strand breaks, base free sites and DNA-protein cross-links [11]. Human body incurs tens of thousands of DNA-damaging events per day [12]. This makes DNA damage a major problem because DNA damage can interfere with cellular processes like transcription or replication and DNA is a repository of genetic information in the cells, and its stability can produce greater consequences than other cellular components such as proteins and RNA.

DNA damage caused by mutations can cause cancer or other diseases, and contribute to the aging process [13]. Thus, cells initiate a highly coordinated cascade of events required for its repair-known as DNA damage response (DDR). Genetically normal cells or individuals have active DNA repair mechanisms, which eventually eliminate almost all damage from DNA. However, some people are born with mutations that affect DNA repair mechanisms, and these people have a high cancer risk [14].

Cells have developed multiple repair mechanisms to repair many types of DNA damage. There are five major DNA repair mechanisms: nucleotide excision repair (NER), mismatch repair (MMR), base excision repair (BER),single strand break repair (SSBs), double-strand break repair (DSBs), which includes both homologous recombination (HR) and non-homologous end joining (NHEJ[15,16]. During the DNA damage response, repair factors are rapidly recruited to the DNA damage site to form a multiprotein repair complex. Damaged DNA is sensed by the repair complex, which activates the DDR.

Researchers have extensively studied PARP-1 (poly (ADP-ribose) polymerase) as it is a well-known regulator of DNA damage repair, especially DNA strand breaks (DSBs) [20]. In addition to this CHK1 is also required for the checkpoint-mediated arrest of cells and the activation of DNA repair in response to DNA damage. The mechanisms by which CHK1 is regulated are complex and comprise multiple steps [24]. CHK1 is activated by its interaction with RAD51 during Homologus recombination, promoting intra-S and G2/M cell cycle checkpoints and modulating the cellular response to replication stress [25].

The ATM (ataxia-telangiectasia mutated) and ATR (ataxia-telangiectasia and Rad3-related) share some overlapping functions. DNA damage sensors activate and recruit ATM, whereas ATR is activated and recruited to DSB sites with its stable binding partner ATRIP (ATR-interacting protein) [18]. ATM and ATR kinases have been considered to be crucial mediators of the DDR due to their ability to promote DDR and mediate cell cycle arrest [19]. CDK1 determines the cell cycle progress by controlling the centrosome cycle, mitotic onset, G2/M transition, G1 progression, and G1/S transition with cyclins [21].

Wee1 assists in the entry of cells into mitosis by negatively regulating the cell cycle entry process. The cyclin B1/CDK1 complex is phosphorylated and inactivated by Wee1 in a specific manner [22]. Wee1 inhibition is considered a promising approach for cancer therapy [23].

There is increasing evidence supporting the use of plant extracts and compounds in the treatment of several malignancies including cancer, neurodegenerative disorders, cardiovascular diseases, and diabetes [26,27]. In addition, they have shown significant potential in modulating the chief signaling pathways involved in cancer progression. Researchers have found that numerous plant-based molecules exhibit significant anticancer effects, including vinblastine, doxorubicin, camptothecin, and paclitaxel [28]. In our previous studies with milk thistle and mushroom, we have shown PARP-1 inhibitors activity [29,30,31].

The antioxidant flavonoid rutin is widely found in natural sources such as fruits (e.g., apples, grapes, lemons), vegetables (e.g., carrots, potatoes), and beverages (e.g., tea and wine). Rutin has been shown to possess a variety of pharmacological properties, including anti-inflammatory, antioxidant, antidiabetic, vasoprotective, antimicrobial, and anticancer properties. Recent studies have shown, in various in vitro cell models, that rutin is able to inhibit tumor growth and induce cell cycle arrest and apoptosis [29].

Naringin is formed from the flavanone naringenin and the disaccharide neohesperidose, and is an active element in Chinese herbal medicine [30]. The compound is widely found in citrus fruits, bergamot, tomatoes, and other fruits, found in glycosides, most notably naringin. This phytochemical has been shown to possess antioxidant, anti-tumor, antiviral, antibacterial, anti-inflammatory, anti-adipogenic, and cardioprotective properties [34].

## Material and Methods

Modified Eagle’s Minimal Essential Medium (DMEM), Phosphate buffer (pH 7.4), trypsin-EDTA, penicillin, streptomycin, Glutamine, fetal bovine serum (FBS), Dimethyl sulfoxide, Trypan blue, Thiazolyl Blue Tetrazolium Bromide were obtained from sigma aldrich (UK). Human Dermal Fibroblasts (2DD) and breast cancer cells (MDA-MB-231) were purchased from Health protection agency culture collection. Disodium phosphate (Na_2_HPO_4_), Monopotassium phosphate (KH_2_PO_4_), Ethylenediaminetetraacetic acid (EDTA) were purchased from sigma Aldrich (UK).

Deoxyribose, Ethylenediaminetetraacetic acid (EDTA), L-ascorbic acid, Iron(III) chloride hexahydrate, Thiobarbituric acid, Trichloroacetic acid Hydrogen peroxide, sodium hydroxide(NaOH), potassium nitrite, Manganese dioxide Diethylenetriamine pentaacetate (DTPA), sodium chloride (NaCl), potassium chloride (KCl), Evans Blue, Nicotinamide adenine dinucleotide, nitro blue tetrazolium, phenazine methosulphate Sodium nitroprusside, sulphanilamide, glacial acetic acid, napthylethylenediamine dihydrochloride (NED) and Griess agent were purchase from sigma Aldrich (UK). All were analytical grades.

### Insilico Studies

*Insilico* study is a computational method to study the chemical compound database to identify molecules with desired biological activity. AutoDock Vina implemented on PyRx 0.8 [35] was used in this study to calculate the binding energies. High throughput Virtual Screening (HTVS) programs through PyRx software with graphical user interfaces (GUIs) that employs AutoDock for predicting receptor–ligand interactions is beneficial for the comparison of ligands (36). AutoDock Vina is a software that works on the premise of empirical scoring functions and also calculates the grid maps automatically.

#### Ligands Selection and Preparation

The structure of biomolecules Naringin and Rutin and standards Olaparib, AZD0156, AD6738, AZD7762, AZD1775 was retrieved by utilizing Pubchem compound database. All the 3D structures of the bioactive molecules were retrieved in structural data format (SDF). The retrieved biomolecules were then minimized using Open Babel using uff force field and conjugate gradients as an optimization algorithm which is available in PyRx 0.8.

#### Preparation of macromolecule

DNA damage response proteins was retrieved from the PDB (https://www.rcsb.org) website and viewed under Discovery Studio 4.0. The retrieved molecules were complex with water molecules and hetero-atoms. Hence, Discovery Studio 4.0 was used to remove hetero atoms and water molecules to avoid docking interference; hydrogen was added and saved in the PDB format. PDB files were retrived from protein data bank for respective targets, including PARP protein (4und), ATM (5np1), ATR (4igk), CHK1 (2yex), WEE1 (7n3u).

#### Ligand protein docking

Docking was performed by using PyRx 0.8. After the completion of docking, autodock preferences were obtained for both ligand and target in PDBQT format. Docking of Protein and ligand were viewed using Discovery studio 4.0 and ligand-protein interaction was analyzed. The pose of minimum binding energy was chosen as the best interaction.

### ADMET and drug-likeness predictions of ligand

The pharmacokinetic properties such as absorption, distribution, metabolism, and excretion and toxicity studies of bioactive molecules play an important role in the drug development steps. Hence, all the possible pharmacokinetic parameters (ADMET) and toxicity of selected biomolecules were predicted using Swiss ADME [37], and PKCSm [38].

### Cell culture

Human dermal fibroblast (2DD) and human breast adenocarcinoma (MDA-MB-231) were grown with the help of culture media.

All ingredients listed in table-1 were pre-warmed at 37^0^C before starting to prepare the medium. All the materials placed in the cabinet were sprayed with bio guard. Hand gloves were used while working in the cabinet to maintain the aseptic conditions.

**Table-1.** Composition of cell culture after modification adopted from (39).

| Sr No | Name of Ingredient | Quantity |
| --- | --- | --- |
| 1 | Dulbecco's Modified Eagle's Medium with a high glucose content (4.5 g l<sup>-1</sup>) | 500 mL |
| 2 | 10% v/v Foetal Bovine serum | 56.5 mL |
| 3 | 100 U ml<sup>-1</sup> Penicillin and 10µg ml<sup>-1</sup> streptomycin | 6mL |
| 4 | 1% Glutamine | 6mL |
| 5 | 1% non-essential amino acids | 6mL |

Ingredients 2-5 from table-1 were measured and added to Dulbecco’s Modified Eagle’s bottle. Medium was filtered using a 0.2µ filter. The cell vials were removed from the nitrogen freezer and placed in a 37^0^C water bath to rapidly defrost the suspension. Cells were plated in 90mm petri dishes and placed into a humidified incubator at 37^0^C with 5% carbon dioxide. Medium was changed on Tuesday and Friday. The cells were passaged twice every week. The medium was removed and the plate with culture was washed using versene (KCl 0.02% (w/v), NaCl 0.8% (w/v), KH2PO4 0.02% (w/v), Na2HPO4 0.0115% (w/v), and 0.2% EDTA (w/v)). The cultures were then treated with a solution of 0.25% trypsin: versene (1:10, v:v) to detach the cells from the tissue culture flasks (approximately 3-5 minutes). The effect of trypsin was neutralized by addition of an equal volume of DMEM medium. This cell suspension was centrifuged at 1000rpm for 5 minutes. The supernatant was removed and the cells were re-suspended in a known volume of fresh medium [40]

#### Cell Counting

Haemocytometer cell counting method was used to count the cells before performing any cell base assays.

### Cytotoxicity assay

#### MTT assay

MTT Assay was performed on the MDA-MB-231 cells to find the cytotoxicity of the molecules on the cells. The molecules were applied in serial dilution. Cytotoxicity is determined by plotting the graph of Cell Viability Vs Concentration.

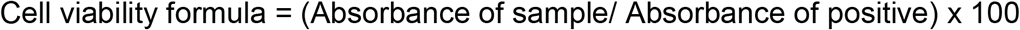

Cells were seeded on 96 well plates at a final concentration of approximately 1.5 × 10^4^ cells per 200µl medium per well 24 hours before the assay. 96 well plates with cell suspensions were then incubated at 37^0^C for 24 hours.

After 24 hours the cell media was removed and the cells were treated with different concentrations of molecules and incubated at 37^0^C for 24 hours.

After 24 hr 20µL MTT (5mg/mL) dye solution in PBS was added to 96 well plates and incubated with cells for 3hrs at 37^0^C. After 3hrs the media containing MTT was removed and the plates were washed with 100 µl of PBS. After washing with PBS the solution of DMSO (200µl) was added to the wells and kept on shaker for 5 to 10 mins. The absorbance was measured at 580nm using a microplate reader [41].

#### DPPH assay

DPPH was performed using a Microplate Reader (BMG BMG LABTECH Instrument). The reaction mixture in each one of the 96-wells consists of molecule solutions, aqueous methanol solution, and 70% ethanol as a blank containing DPPH radicals. The mixture was left to stand for 60 min in the dark. The reduction of the DPPH radical was determined by measuring the absorption at 517 nm [42].

#### Hydroxyl radical scavenging activity assay

Assay was adopted from [43] with slight modification. Test tubes were used to perform the procedure which was then transferred to a 96 well plate to read the absorbance in 96 well plate.

First a mixture was prepared in which 3.6 mM deoxyribose, 0.1 mM EDTA, 0.1 mM L-ascorbic acid, 1 mM H2O2 and 0.1 mM Iron(III) chloride hexahydrate (FeCl3.6H2O) are added and then 10µM of molecules were added to this mixture, and volume was made up to 1mL with 25 mM phosphate buffer, pH 7.4. This mixture was incubated for 1hr at 37°C.

After one hour 500 μl of 1% Thiobarbituric acid and 500 μl of 1% Trichloroacetic acid were added to the mixture and then heated in a water-bath (80°C) for 20 min and then cooled. The absorbance was measured at 532 nm. The control reaction contained no test sample.

#### Peroxynitrite scavenging activity assay

Peroxynitrite assay is divided into two parts. First peroxynitrite is produced and in the next step fractions are applied. Peroxynitrite assay is divided in two part. First Peroxynitrite were synthesis according to and then evans blue bleaching assay was used to measure the peroxynitrite scavenging activity [40]

In the first Step, an acidic solution of 0.7 M H_2_O_2_ was mixed with an equal volume of 0.6 M potassium nitrite in an ice bath and an equal volume of ice cold 1.2 M NaOH was added. Granular Manganese dioxide prewashed with 1.2 M NaOH was used to remove excess H_2_O_2_ and the reaction mixture was left at −20°C for 12hrs.

In the second step, Evans blue bleaching is performed. The reaction mixture consisted of 0.1 mM DTPA, 90 mM NaCl, 5 mM KCl, 12.5 μM Evans Blue and molecules (10µM) were added to first step produced peroxynitrite and final volume was adjusted to 1 ml with 50 mM phosphate buffer (pH 7.4). The reaction mixture was incubated at 25°C for 30 min and the absorbance was measured at 611 nm. The percentage of scavenging of ONOO-was calculated by comparing the results of the test and blank samples.

#### Superoxide radical scavenging activity assay

Superoxide radical scavenging activity was measured by using non-enzymatic involving the nicotinamide adenine dinucleotide-nitro blue tetrazolium-phenazine methosulphate (NADH-NBT-PMS) system as reported by [44].

The superoxide radical’s scavengers were assayed by inhibition of NBT reduction by NADH in the presence of PMS, which reduce nitro blue tetrazolium (NBT) to a purple formazan. NBT (50 μM in 20 mM phosphate buffer, pH 7.4) was added to 1 ml of NADH solution (73 μM of NADH in 20 mM phosphate buffer, pH 7.4) in the presence of 10µM of molecules. The reactions were initiated by adding PMS (15 μM) and the absorbance was measured at 560 nm. The percentage of scavenging of superioxide radical scavenging was calculated by comparing the results of the test and blank samples.

#### Nitric oxide scavenging activity assay

Nitric oxide is generated at physiological temperature from aqueous sodium nitroprusside (SNP) solution after reacting with oxygen to produce nitrite ions, which are detected by Griess Illosvoy reaction [45].

The reaction mixture contained 10 mM SNP in 20 mM phosphate buffer, pH 7.4 and 10µM of molecules in a final volume of 3 ml. After incubation for 150 min at 25°C, 1 ml of sulfanilamide (0.33% in 20% glacial acetic acid) was added to 0.5 ml of the incubated solution and allowed to stand for 5 min. Then 1 ml of NED (0.1% w/v) was added and the mixture was incubated for 30 min at 25°C. The absorbance was measured at 540 nm. The percentage of scavenging of superoxide radical scavenging was calculated by comparing the results of the test and blank samples. In this assay pink chromophor generated during diazotization of nitrite ions with sulphanilamide and subsequent coupling with NED was measured.

#### Single Cell gel electrophoresis (COMET) assay

Cells were grown in small petri dishes with normal cell culture protocol for a week. 2DD cells were grown in low serum media (0.5%) was added to the dishes and left for 7 days to make them quiescent fibroblast cells (QFC). MDA-MB-231 cells were grown using the normal protocol as mentioned previously. On the experiment day the media was removed and cells were washed twice with PBS. The cells were treated with molecules for one hour. After one hour cells were washed twice with pre-warmed PBS. Trypsin-EDTA was added to coat the entire monolayer of cells. Cells are incubated for 2 minutes at 37° C or until cells easily detached upon tapping. 2 mL of complete media (containing fetal bovine serum) was added to inactivate the trypsin. Cells are transferred to a centrifuge tube and re-suspended at 1.5 × 10^5^ cells/mL in ice cold 1X PBS.

The comet assay involves lysis with detergent and high salt after embedding cells in agarose so that the DNA is immobilized for subsequent electrophoresis. Slides are dried overnight at 37° C. Drying brings all the cells to a single plane which facilitates observation. Samples may be stored at room temperature with desiccant prior to scoring. 100µL of diluted SYBR Green I was added onto each circle of dried agarose and placed in the refrigerator for 15 - 30 minutes.

The assay was also adapted to detect Oxidative base damage by adding enzyme Formamidopyrimidine DNA glycosylase (FPG) which combines a specific glycosylase activity, removing the damaged base and creating an apurinic/apyrimidinic (AP) site, and an AP lyase which converts the AP site to a break[40].

## Result and Discussion

*Insilico* studies were performed using PyRx to identify the binding affinity between proteins and the molecules. Here we used PARP-1, ATM, ATR, CHK1 and WEE1 proteins from DNA damage response pathway. The results are mentioned in table-2.

The result suggests that Naringin has shown good binding affinity against ATM, ATR, CHK1 and WEE1 while Rutin has shown good binding affinity against PARP-1,ATM and ATR.

**Table-2.**
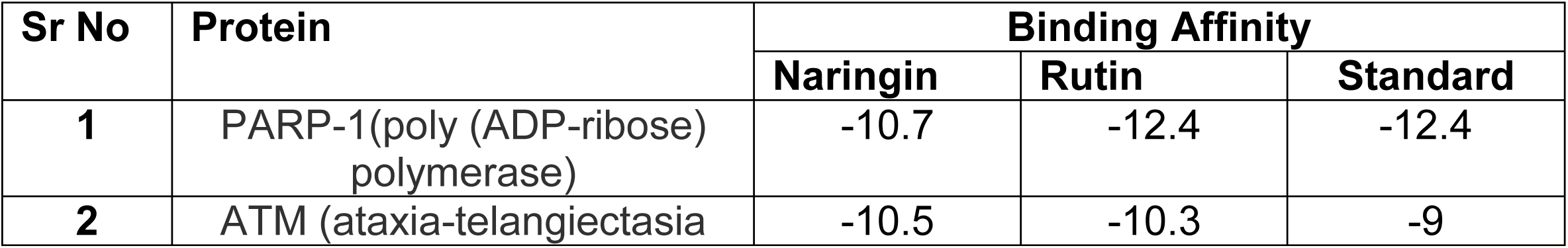
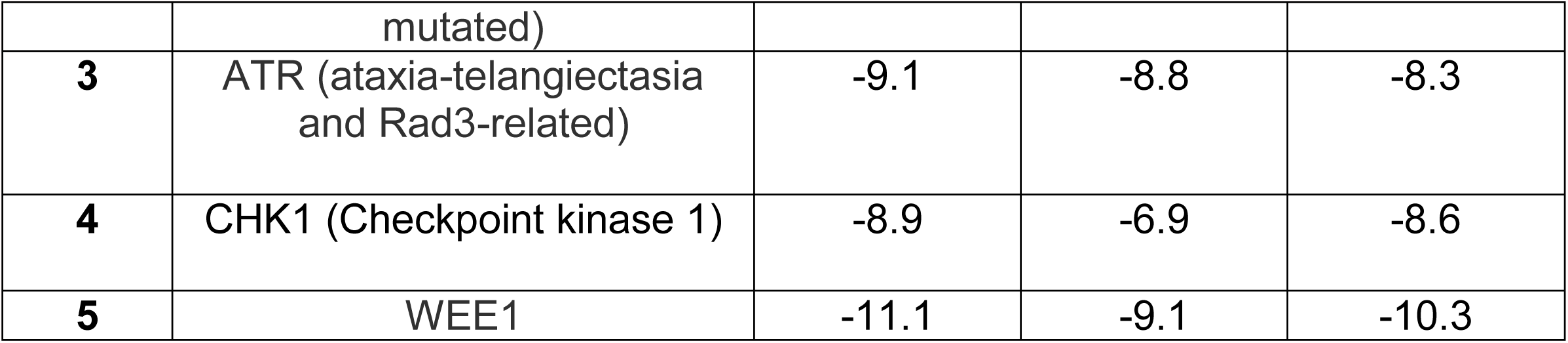
Analysis of DDR protein binding affinity with Naringin and Rutin.

| Sr No | Protein | Binding Affinity |  |  |
| --- | --- | --- | --- | --- |
|  |  | Naringin | Rutin | Standard |
| 1 | PARP-1(poly (ADP-ribose) polymerase) | -10.7 | -12.4 | -12.4 |
| 2 | ATM (ataxia-telangiectasia mutated) | -10.5 | -10.3 | -9 |
| **3** | ATR (ataxia-telangiectasia and Rad3-related) | -9.1 | -8.8 | -8.3 |
| **4** | CHK1 (Checkpoint kinase 1) | -8.9 | -6.9 | -8.6 |
| **5** | WEE1 | -11.1 | -9.1 | -10.3 |

**Table-3.** ADMET SWISS profile of Naringin and Rutin.

| Sr No | Properties | Naringin | Rutin |
| --- | --- | --- | --- |
| **Physicochemical Properties** |  |  |  |
| 1 | Formula | C<sub>27</sub>H<sub>32</sub>O<sub>14</sub> | C<sub>27</sub>H<sub>30</sub>O<sub>16</sub> |
| 2 | Molecular weight | 580.53 g/mol | 610.52 g/mol |
| 3 | Num. heavy atoms | 41 | 43 |
| 4 | Num. arom. heavy atoms | 12 | 16 |
| 5 | Fraction Csp<sup>3</sup> | 0.52 | 0.44 |
| 6 | Num. rotatable bonds | 6 | 6 |
| 7 | Num. H-bond acceptors | 14 | 16 |
| 8 | Num. H-bond donors | 8 | 10 |
| 9 | Molar Refractivity | 134.91 | 141.38 |
| 10 | TPSA | 225.06 Å<sup>2</sup> | 269.43 Å<sup>2</sup> |
| **Lipophilicity** |  |  |  |
| 11 | Log *P*<sub>o/w</sub> (iLOGP) | 2.38 | 2.43 |
| 12 | Log *P*<sub>o/w</sub> (XLOGP3) | -0.44 | -0.33 |
| 13 | Log *P*<sub>o/w</sub> (WLOGP) | -1.49 | -1.69 |
| 14 | Log *P*<sub>o/w</sub> (MLOGP) | -2.77 | -3.89 |
| 15 | Log *P*<sub>o/w</sub> (SILICOS-IT) | -1.64 | -2.11 |
| 16 | Consensus Log *P*<sub>o/w</sub> | -0.79 | -1.12 |
| **Water Solubility** |  |  |  |
| 17 | Log S (ESOL) | -2.98 | -3.30 |
| 18 | Solubility | 6.04e-01 mg/ml ;<br>1.04e-03 mol/l | 3.08e-01 mg/ml ; 5.05e-04 mol/l |
| 19 | Class | Soluble | Soluble |
| 20 | Log S (Ali) | -3.82 | -4.87 |
| 21 | Solubility | 8.77e-02 mg/ml ;<br>1.51e-04 mol/l | 8.30e-03 mg/ml ; 1.36e-05 mol/l |
| 22 | Class | Soluble | Moderately soluble |
| 23 | Log S (SILICOS-IT) | -0.49 | -0.29 |
| 24 | Solubility | 1.87e+02 mg/ml ;<br>3.21e-01 mol/l | 3.15e+02 mg/ml ; 5.15e-01 mol/l |
| 25 | Class | Soluble | Soluble |
| **Pharmacokinetics** |  |  |  |
| 26 | GI absorption | Low | Low |
| 27 | BBB permeant | No | No |
| 28 | P-gp substrate | Yes | Yes |
| 29 | CYP1A2 inhibitor | No | No |
| 30 | CYP2C19 inhibitor | No | No |
| 31 | CYP2C9 inhibitor | No | No |
| 32 | CYP2D6 inhibitor | No | No |
| 33 | CYP3A4 inhibitor | No | No |
| 34 | Log $K_p$ (skin permeation) | -10.15 cm/s | -10.26 cm/s |
| **Druglikeness** |  |  |  |
| 35 | Lipinski | No; 3 violations:<br>MW>500,<br>NorO>10,<br>NHorOH>5 | No; 3 violations:<br>MW>500,<br>NorO>10,<br>NHorOH>5 |
| 36 | Ghose | No; 4 violations:<br>MW>480,<br>WLOGP<-0.4,<br>MR>130,<br>#atoms>70 | No; 4 violations:<br>MW>480,<br>WLOGP<-0.4, MR>130,<br>#atoms>70 |
| 37 | Veber | No; 1 violation:<br>TPSA>140 | No; 1 violation:<br>TPSA>140 |
| 38 | Egan | No; 1 violation:<br>TPSA>131.6 | No; 1 violation:<br>TPSA>131.6 |
| 39 | Muegge | No; 3 violations:<br>TPSA>150, H-acc>10, H-don>5 | No; 4 violations:<br>MW>600, TPSA>150, H-acc>10, H-don>5 |
| 40 | Bioavailability Score | 0.17 | 0.17 |
| **Medicinal Chemistry** |  |  |  |
| 41 | PAINS | 0 alert | 1 alert: catechol_A |
| 42 | Brenk | 0 alert | 1 alert: catechol |
| 43 | Leadlikeness | No; 1 violation:<br>MW>350 | No; 1 violation: MW>350 |
| 44 | Synthetic accessibility | 6.16 | 6.52 |

**Table-4.**
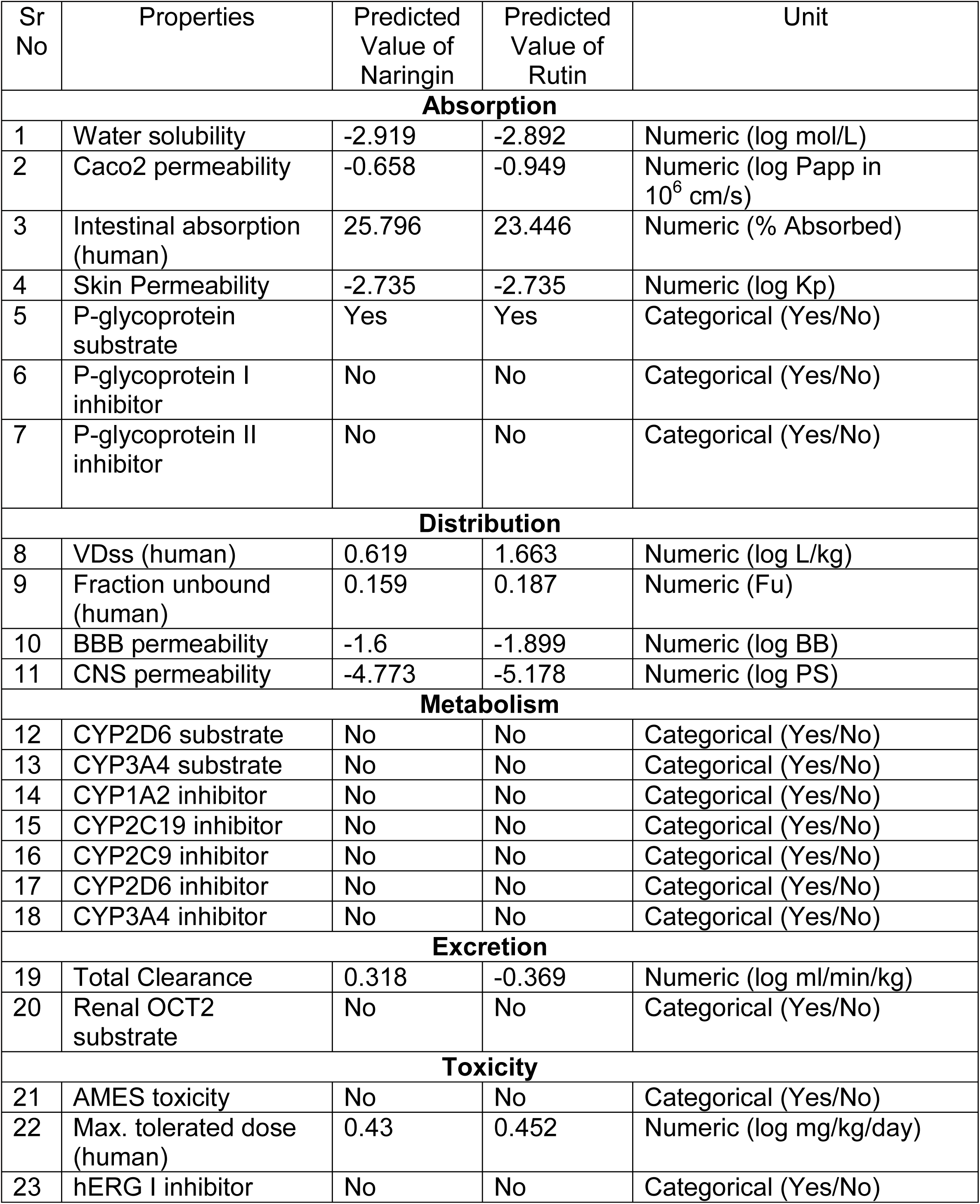
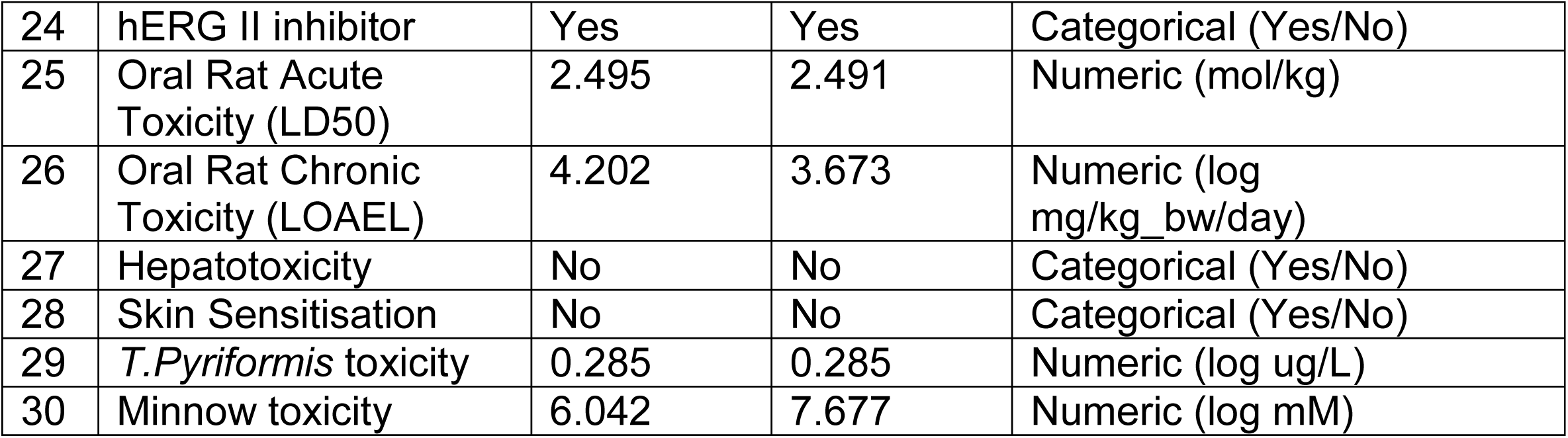
ADMET PkCSM profile of Naringin and Rutin.

The above figure shows the 3D interactions of different standards with DNA damage response pathway proteins. The 3D structure helps to compare the results with the 3D interaction of the molecules.

Figure 2 shows the 3D and 2D interaction of rutin with PARP-1 that help us to understand the binding of molecules at the active site

**Fig-1.**
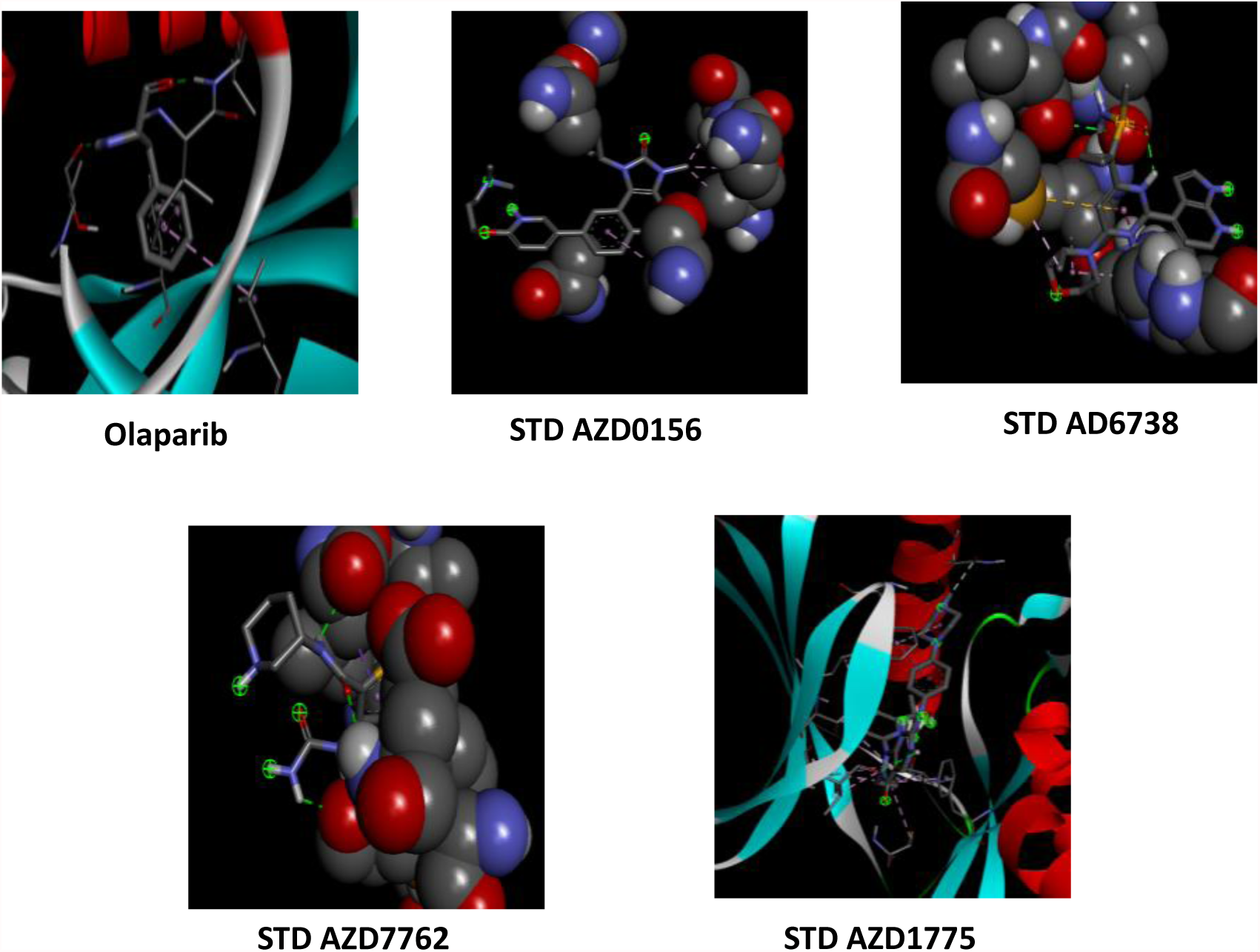
3D structure of standards used in Pyrx based molecules docking.

**Fig-2.**
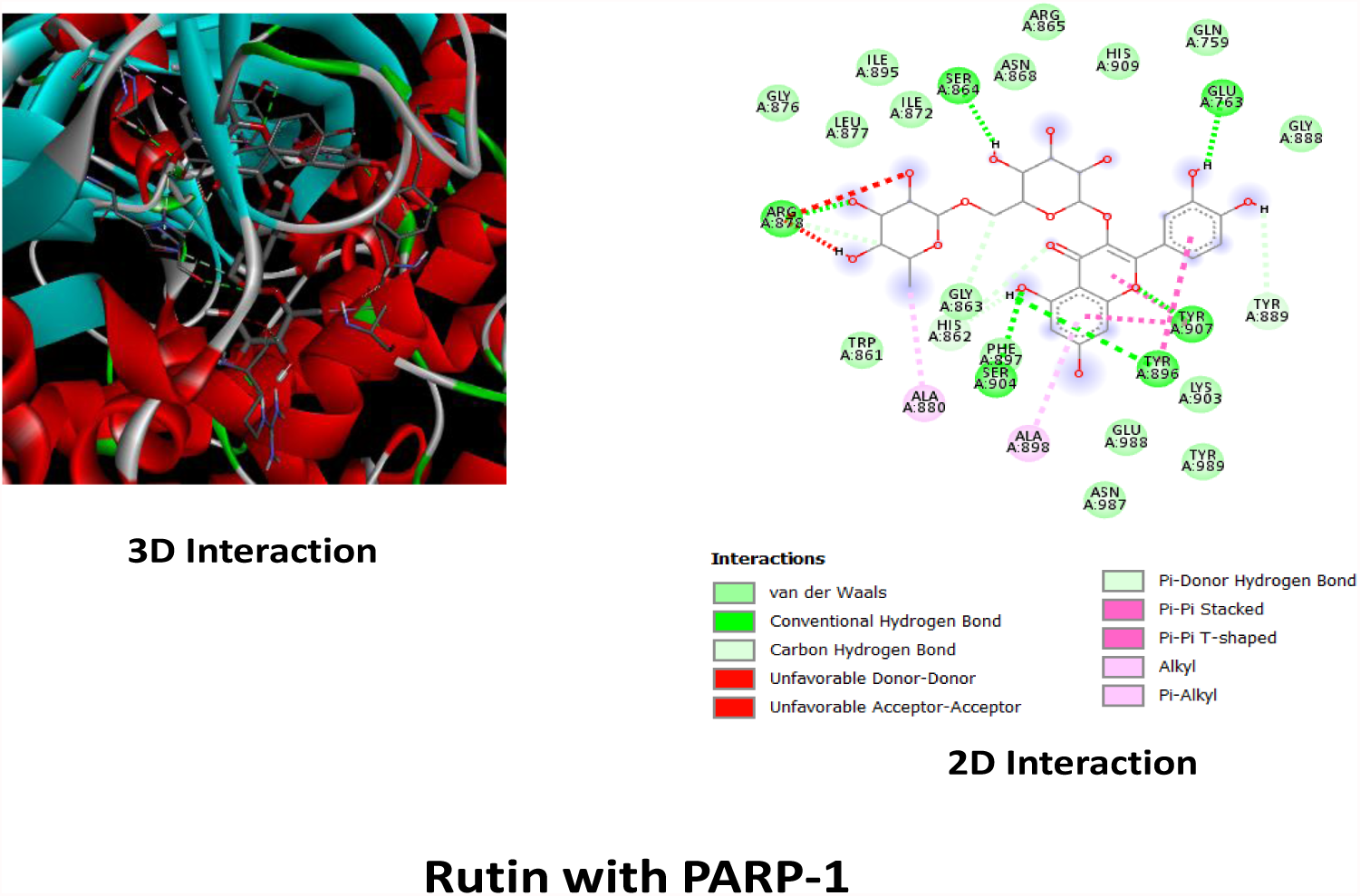
3D and 2D interaction of Rutin with PARP-1

The two-dimensional interaction of ligand rutin with receptor PARP-1 visualized in Discovery Studio showing the residues and type of interactions formed, the ligand formed 6 conventional hydrogen bond with SER864, ARG878,SER904,TYR896, TYR907,GLU763 represented in green It formed hydrophobic bonds with ALA880, ALA898 represented in pink colour. Three-dimensional interaction of rutin with PARP 1 protein visualized using Discovery Studio showing the interacting in dotted line.

Figure 3 shows the 3D and 2D interaction of Naringin and Rutin with ATM that help us to understand the binding of molecules at the active site. The 2D interaction shows the bonds between the group from Naringin and rutin with ATM.

**Fig-3.**
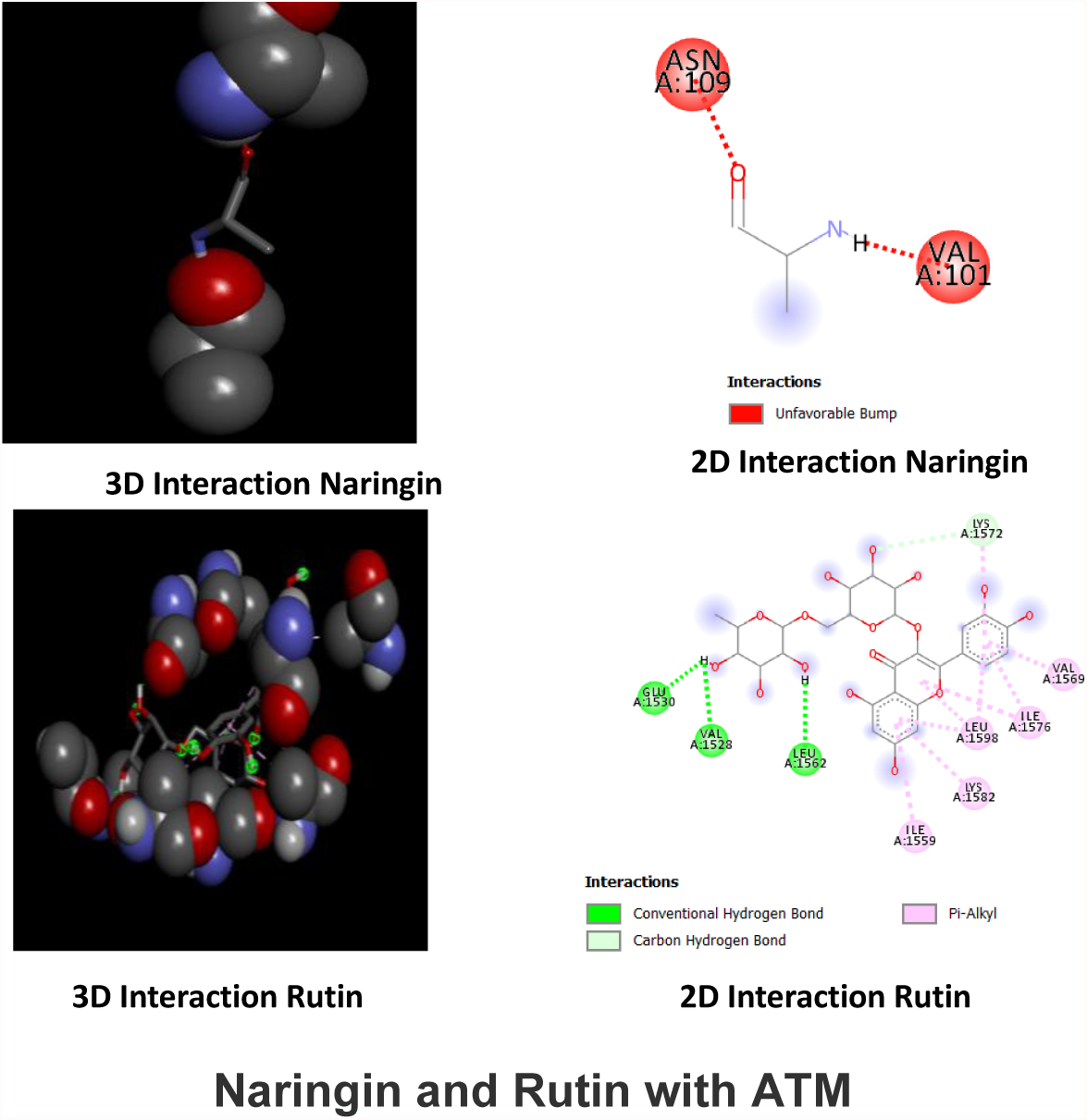
3D and 2D interaction of Naringin and Rutin with ATM.

The two-dimensional interaction of ligand naringin with receptor ATM visualized in Discovery Studio showing the residues and type of interactions formed, the ligand formed unfavourable bump with ASN109, VAL101 represented in red colour. Three-dimensional interaction of rutin with ATM protein visualized using Discovery Studio showing the interacting red line.

The two-dimensional interaction of ligand rutin with receptor ATM showing the residues and type of interactions formed, rutin formed 3 conventional hydrogen bond with GLU1530, VAL1528,LEU1562 represented in green colour.It has formed 5 PiAlkyl bonds with ILE1559, LYS1582,ILE1576,VAL1569, represented in pink colour.It has formed carbon hydrogen bond with LYS1572.Three-dimensional interaction of rutin with ATM protein showing the interacting in green circles, dotted lines.

Figure 4 shows the 3D and 2D interaction of Naringin and Rutin with ATR that help us to understand the binding of molecules at the active site. The 2D interaction shows the bonds between the group from Naringin and rutin with ATR.

**Fig-4.**
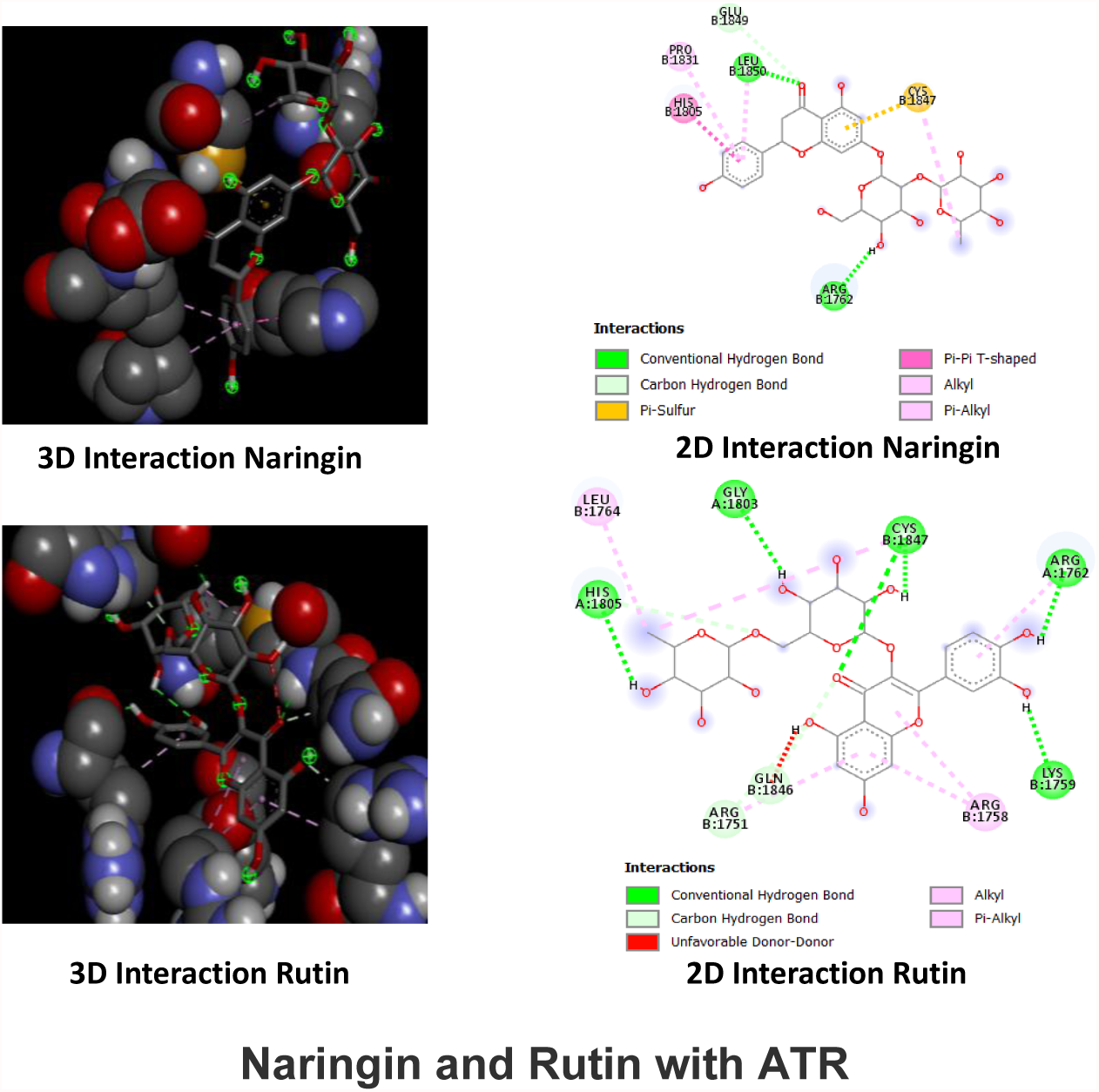
3D and 2D interaction of Naringin and Rutin with ATR.

The two-dimensional interaction of ligand naringin with receptor ATR visualized in Discovery Studio showing the residues and type of interactions formed. The ligand formed 2 conventional hydrogen bonds with LEU1850, ARG1762 represented in green colour. Three-dimensional interaction of naringin with ATR protein visualized using Discovery Studio showing the interaction in green circles and dotted lines.

The two-dimensional interaction of ligand rutin with receptor ATR showing the residues and type of interactions formed. Rutin formed 5 conventional hydrogen bonds with HIS1805,GLY1803,CYS1847,ARG1762,LYS1759 represented in green colour and alkyl and PiAlkyl interaction with ARG1758,LEU1764 represented in pink colour. It has formed a carbon hydrogen bond with GLN1846, ARG1751. Three-dimensional interaction of rutin with ATR protein showing the interaction in green circles, dotted lines.

Figure 5 shows the 3D and 2D interaction of Naringin with CHK1 that help us to understand the binding of molecules at the active site. The 2D interaction shows the bonds between the group from Naringin and CHK1.

**Fig-5.**
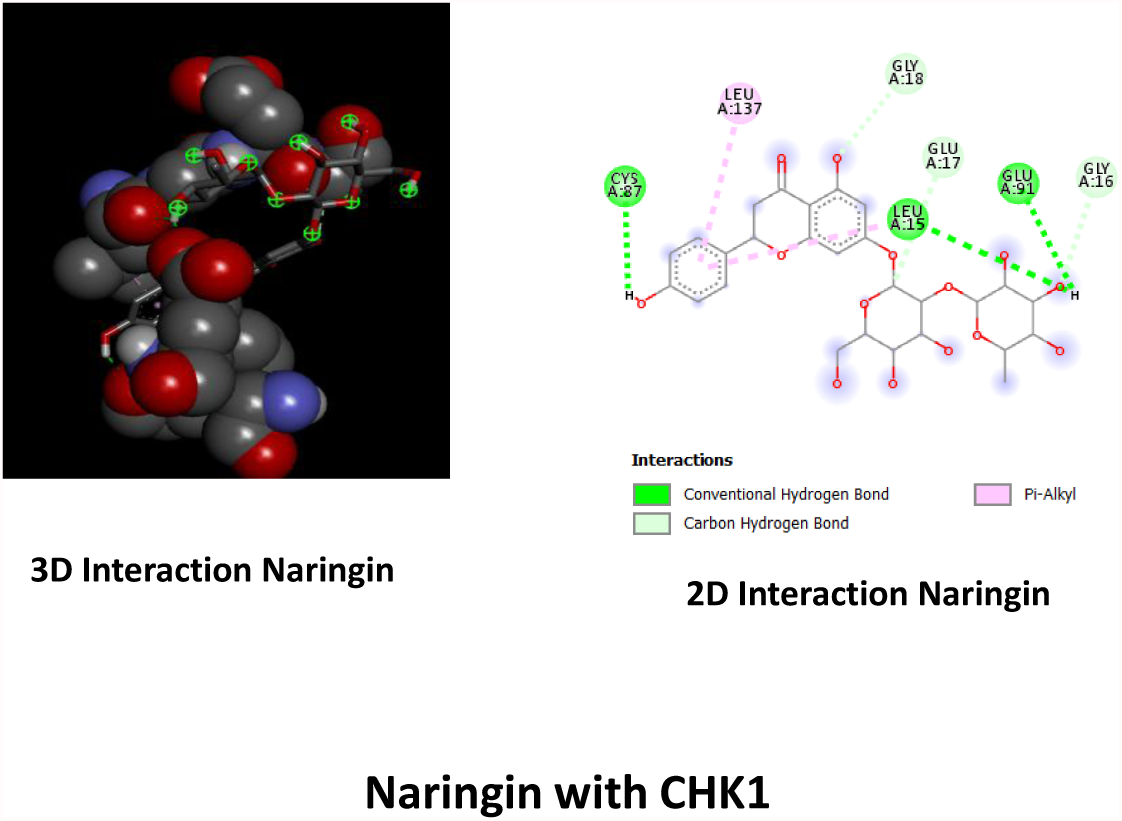
3D and 2D interaction of Naringin with CHK1.

The two-dimensional interaction of ligand naringin with CHK1 visualized in Discovery Studio showing the residues and type of interactions formed, the ligand formed 3 conventional hydrogen bonds with CYS87,LUE15,GLU91 represented in green colour and PiAlkyl interaction with LEU137. It has formed carbon hydrogen bond with GLY18,GLU17, GLY16.

Three-dimensional interaction of naringin with CHK1 visualized using Discovery Studio showing the interacting in green circles and dotted lines.

Figure 6 shows the 3D and 2D interaction of Naringin with WEE1 that help us to understand the binding of molecules at the active site. The 2D interaction shows the bonds between the group from Naringin and WEE1.

**Fig-6.**
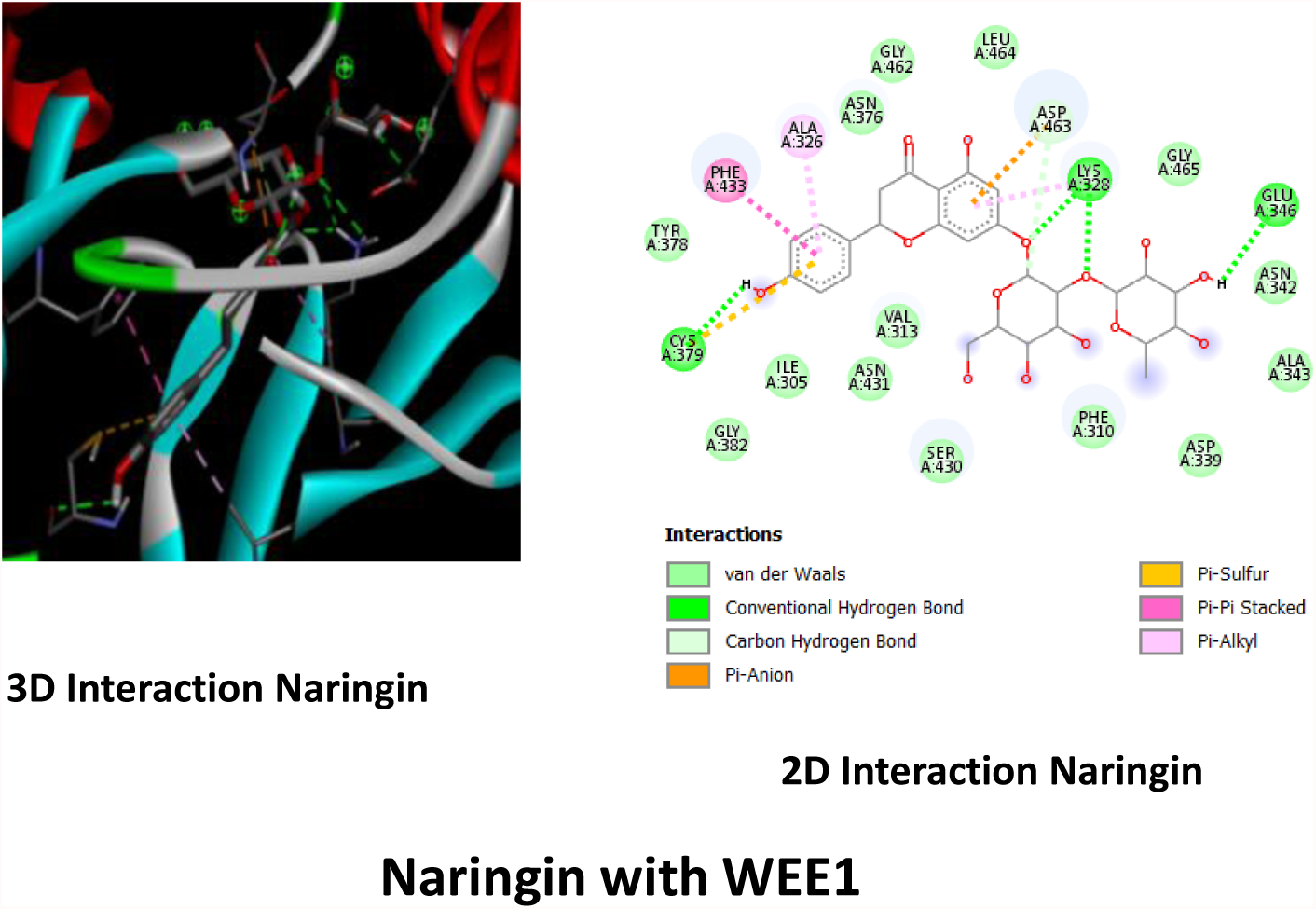
3D and 2D interaction of Naringin with WEE1.

**Fig-7.**
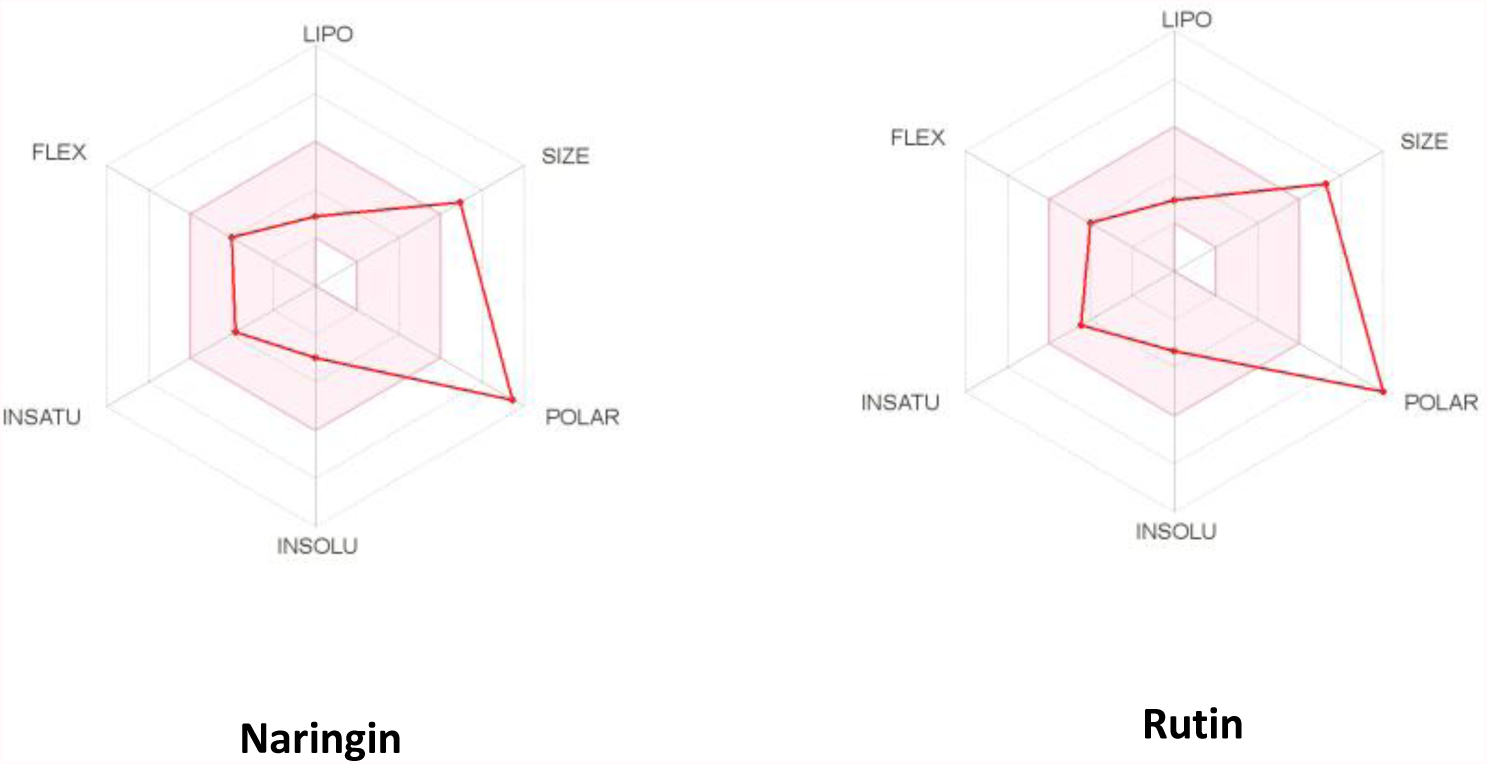
Naringin and Rutin SWISS profile

**Fig-8.**
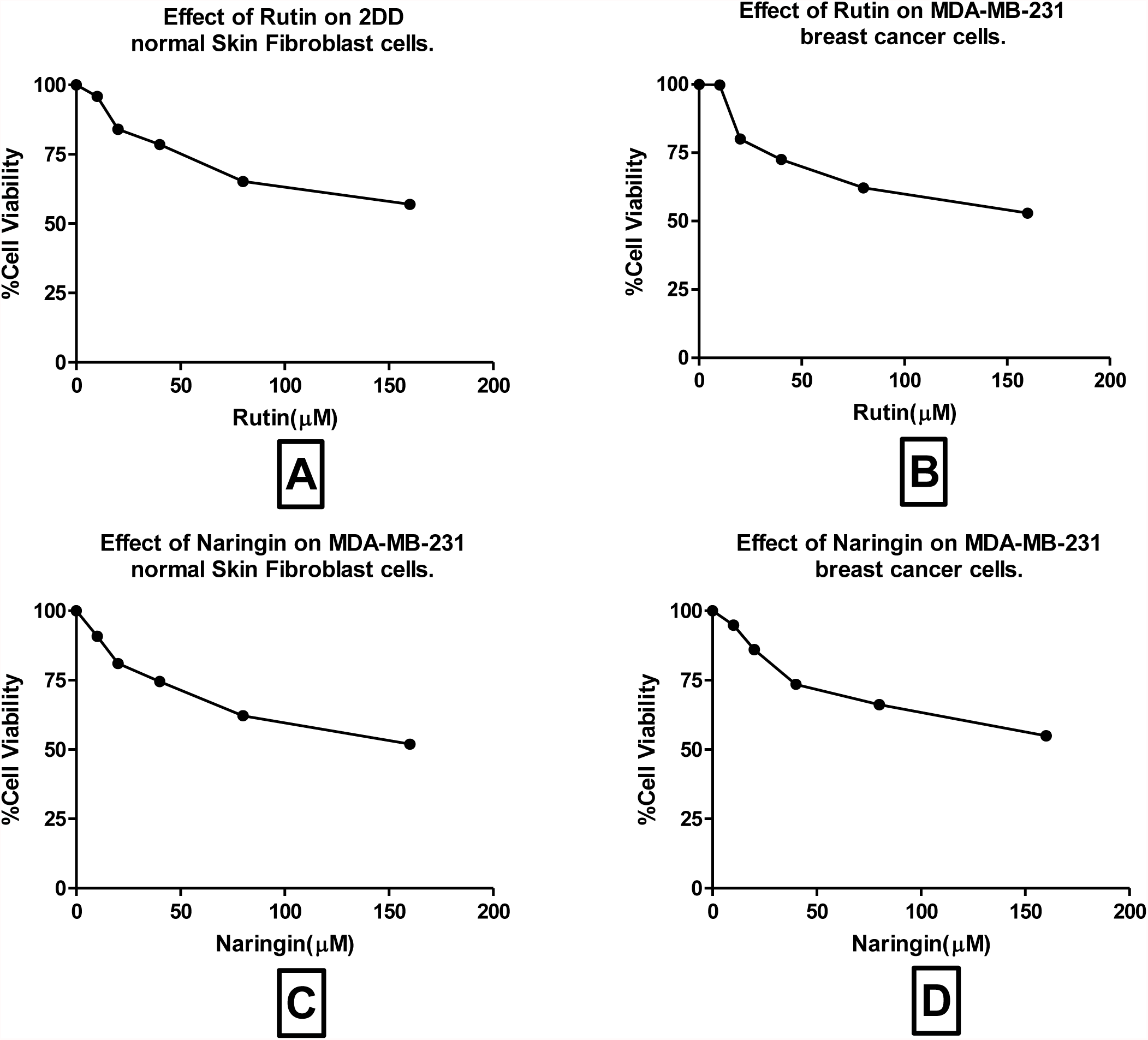
MTT assays of Rutin and Naringin compounds were applied to 2DD nomal skin fibroblast cells and MDA-MB-231 breast cancer cells to identify its cytotoxicity on cells. Molecules were applied in different concentrations (0-160µM).

**Fig-9.**
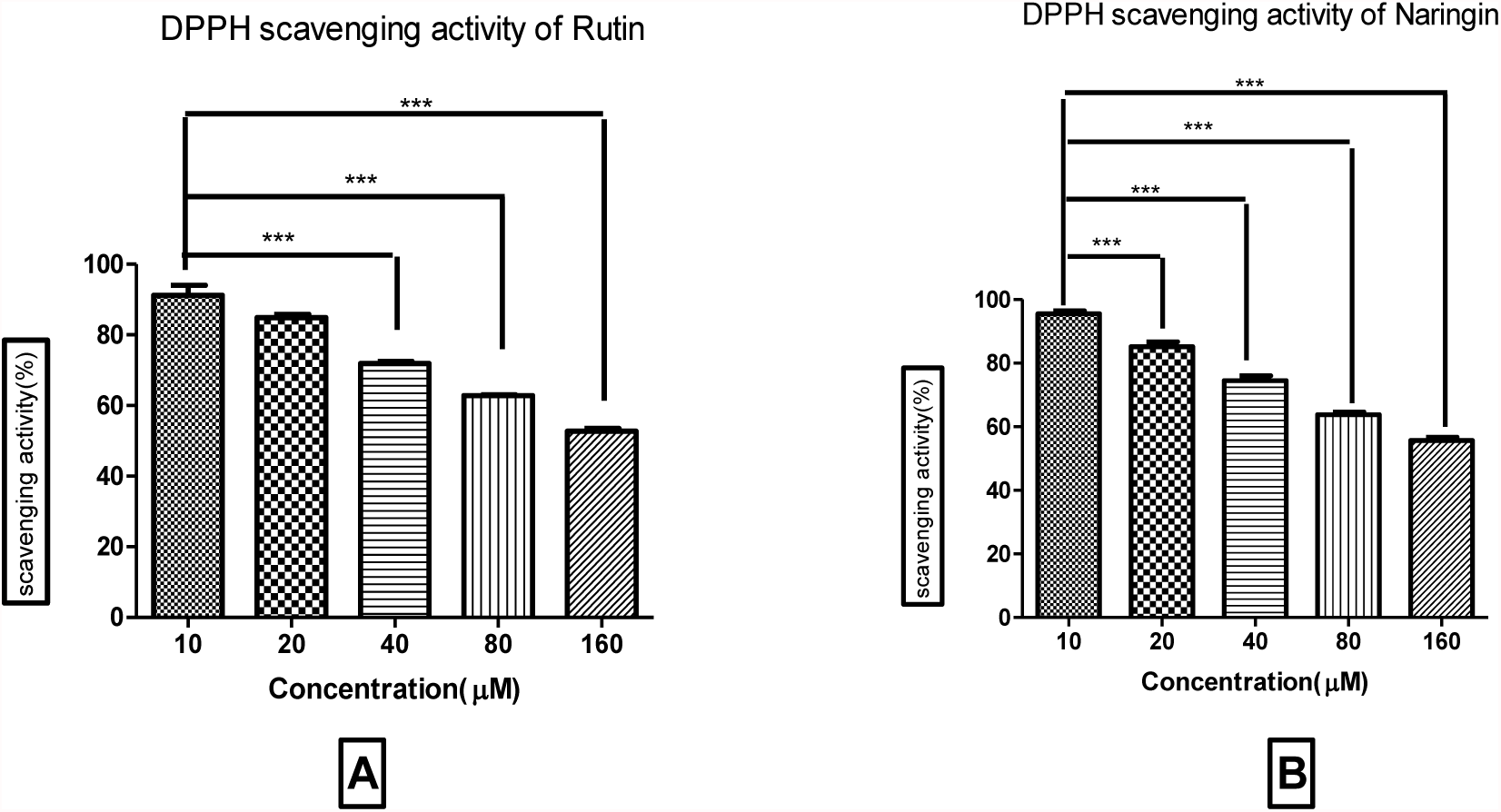
DPPH assay of Rutin(A) and Naringin(A) was tested to identify the free radical scavenging activity. P value was less than 0.0001. Anova was performed followed by Tukey test.

**Fig-10.**
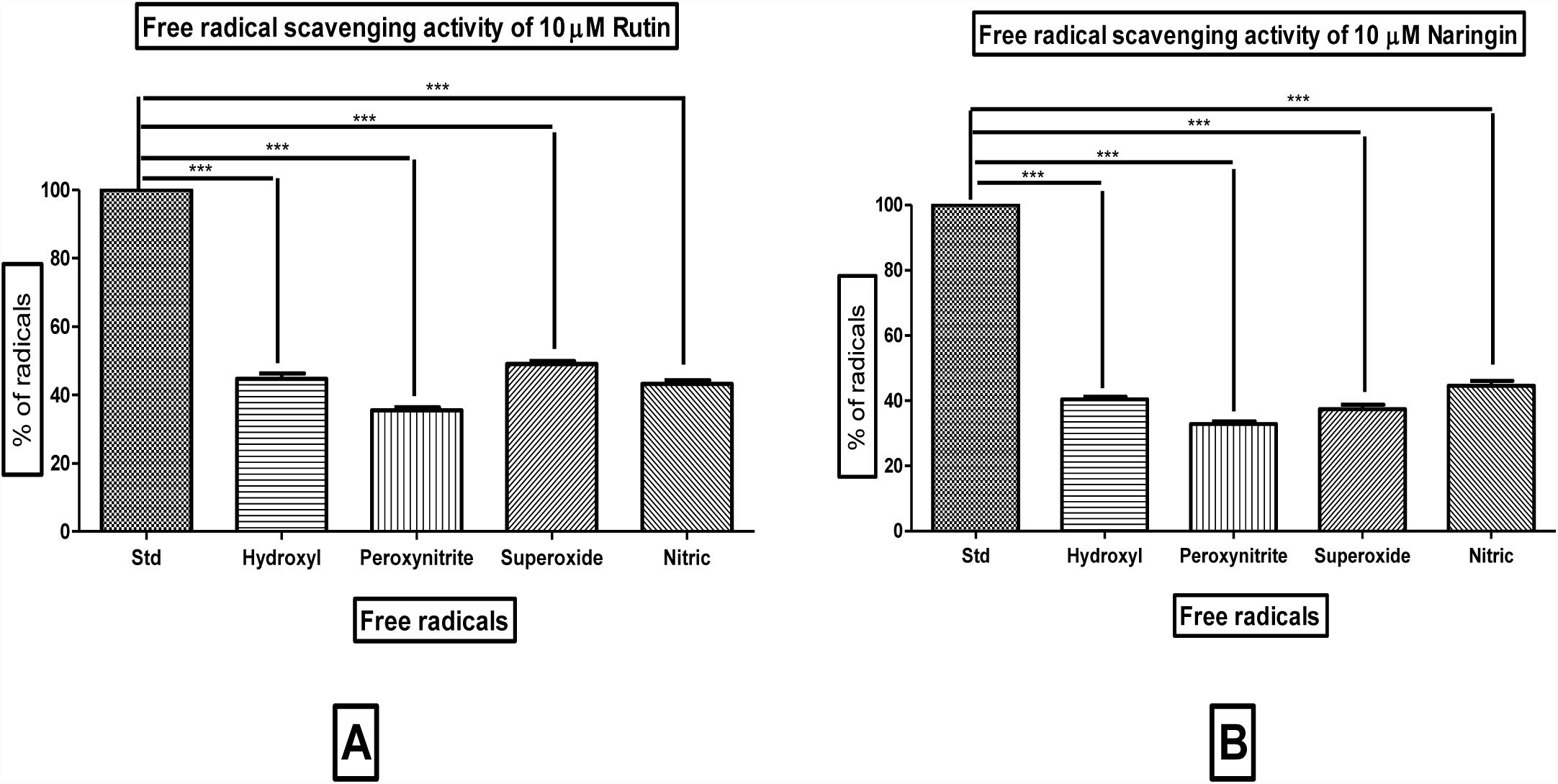
Free radical assay to identify the effect of Rutin (A) and Narigin (B) on Hydroxyl radical, peroxynitrite, superoxide and nitric free radicals. Concentration used was 10 µM, P value was < 0.0001. Anova was performed followed by Tukey Test.

**Fig-11.**
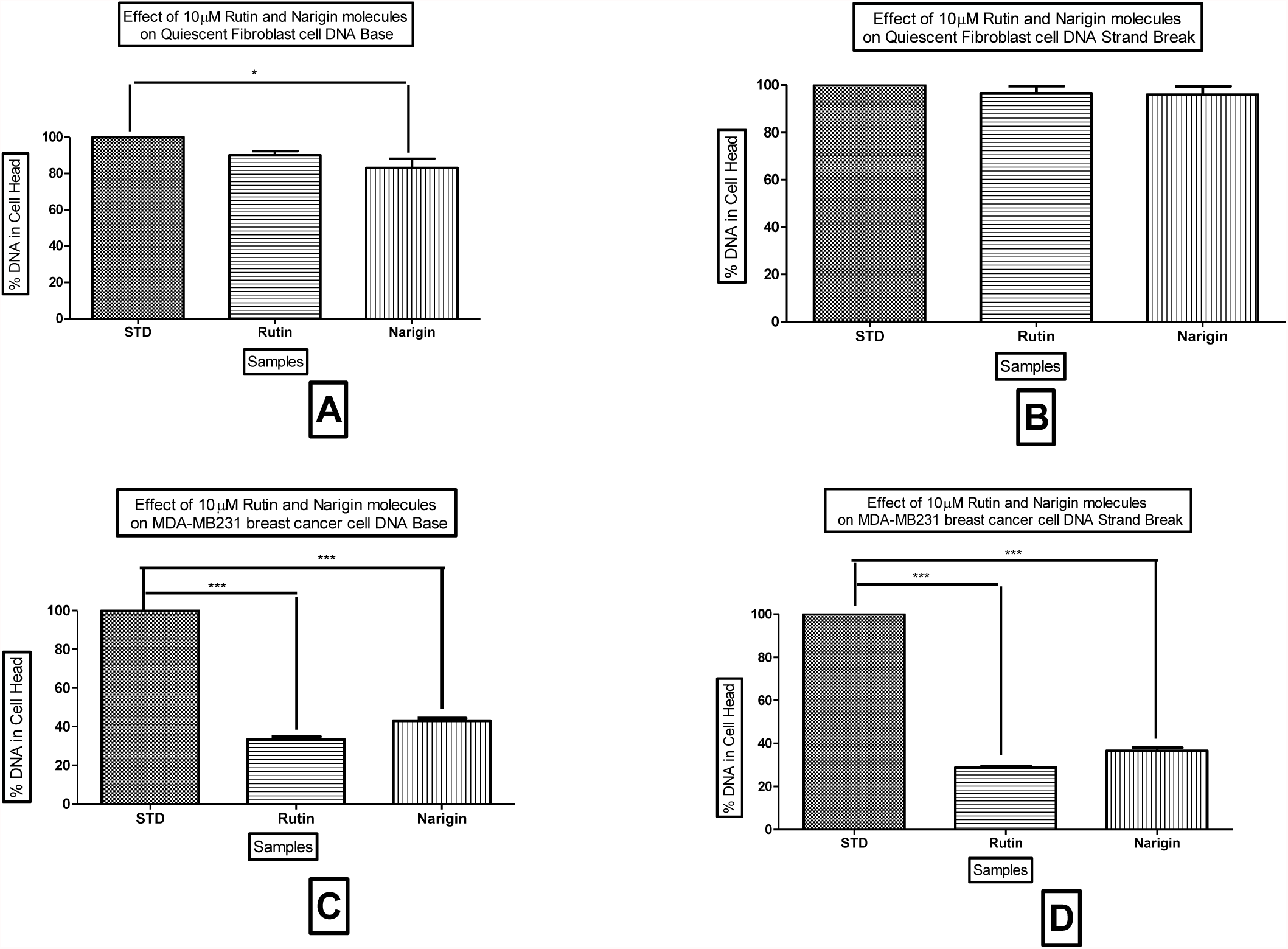
**A**- Single cell gel electrophoresis (COMET assay) of Naringin and Rutin performed on quiescent Fibroblast cell(QFC) to study DNA base damage, **B**- Comet assay of Naringin and Rutin performed on quiescent Fibroblast cell(QFC) to study DNA strand break, **C**- COMET assay of Naringin and Rutin performed on MDA-MB-231 breast cancer cells to study DNA base damage, **D**- Comet assay of Naringin and Rutin performed on MDA-MB-231 breast cancer cells to study DNA strand break. Molecules were applied at 10µM. Each experiment was done in triplicate. Data were represented as means ± Standard Derivation. P value was < 0.0001. Anova was performed followed by Tukey Test.

The two-dimensional interaction of ligand naringin with WEE1 visualized in Discovery Studio showing the residues and type of interactions formed, the ligand formed 3 conventional hydrogen bonds with CYS379,LYS328,GLU346 represented in green colour. It has formed PiAlkyl and Pi-Pi stacked interaction with ALA326 and PHE433. It has formed carbon hydrogen bond with ASP463.

Three-dimensional interaction of naringin with WEE1 visualized using Discovery Studio showing the interacting in green circles and dotted lines.

### ADMET and drug-likeness predictions of ligand

#### Swiss ADME profile

SWISS ADMET helps to understand the profile of Naringin and Rutin. ADMET of the molecules are predicted and help to understand different properties such as Physicochemical Properties, Lipophilicity, Water Solubility, Pharmacokinetics and Druglikeness.

We also used PkCSM ADMET to confirm the profile of Naringin and Rutin. PkCSM ADMET helps us to understand Absorption, Distribution, Metabolism, Excretion and Toxicity.

ADMET and druglikeness data from insilico is helpful to understand the lead at initial stage and further optimization of the molecules can be done invitro.

### MTT assay

Each experiment was done in triplicate. Data were represented as means ± Standard Derivation.

Molecules were tested with MTT assay in 2DD and MDA-MB-231 cells. Percentage of cell viability was observed to be in the range of 75-95% at applied concentration.

MTT assay was performed to identify the concentration required for molecules to produce toxicity in normal and cancer cells. The data suggest that the concentration applied (0-160µM) didnt produce the toxicity.

### DPPH Assay

Each experiment was done in triplicate. Data were represented as means ± Standard Derivation. Statistical analysis was performed by Anova followed by post-hoc tukey test. Tukey test compares std with different concentrations and the stars indicate a significant difference (^***^P<0.0001) between free radicals treated with different concentrations of molecules.

Molecules were screened at the same concentration used in MTT assay. Significant antioxidant activity was observed at a low concentration (10µM).

Based on the MTT and DPPH assay we selected 10µM concentrations of Naringin and Rutin to identify their free radical scavenging activity.

The molecules showed reduced OH, ONOO, O2− and NO free radical scavenging activity. The free radical scavenging is observed more than 60% in naringin applied assays.

Free radical assay data confirm the antioxidant activity of Narigin and rutin. The molecules were further tested for genotoxicity to understand it effect of DNA strand break and base break at 10µM

### Rutin and Naringin effect on DNA base and DNA strand break in Normal and Cancer cells

DNA in the cell head was analysed in 200 cells. Cells were treated with 10µM of molecules. Results expressed as percentage of DNA in cell head in STD=without molecules, Rutin and Naringin.

DNA damage response (DDR) is a complex network of signaling pathways developed by cells to deal with DNA damage. DNA integrity is constantly monitored by DDR, which activates transient cell cycle arrest and DNA repair in the event of any abnormality of DNA that threatens the integrity of DNA [1,2]. Cancer and DNA damage response integrity are not only strictly related, it can also be regarded as the Achilles heel of tumors. At many stages of cancer development, functional inactivation of DDR pathways is a hallmark of the disease.

The accumulation of genetic lesions and the increase in genomic instability contribute to carcinogenesis. However, defects in DDR may reduce the amount of DDR activity remaining in cancer cells, making them more vulnerable to therapy [46]

Various DDR pathways have been characterized, making them attractive targets for tissue-specific inhibitors in the fight against cancer. It is possible to exploit this inhibition by sensitizing tumor cells to the effects of standard genotoxic treatments. A DDR defect in a tumor may be a targetable weakness exploiting this concept of synthetic lethality. Cancer cells may be harmed by targeting DDR pathways that remain intact. Several small molecule inhibitors of poly(ADP-ribose)-polymerase (PARP) have been demonstrated to be highly effective in the treatment of cells harboring mutations in the breast and ovarian cancer susceptibility genes BRCA1 and BRCA2 [47,48]. Further, drug combinations that target more than one non-redundant DDR component may be therapeutically effective.

The focus of this study is on the proteins ATM, ATR, CHK1 and WEE1, discussing their roles in regulating DNA damage response and potential uses as cancer therapy targets. These proteins have been targeted by specific inhibitors for over a decade, which have been investigated for possible anticancer activity as monotherapy strategies against tumors with specific defects (a synthetic lethality approach) as well as in combination with radiotherapy and chemotherapeutics. These inhibitors’ antitumor activity will be critically evaluated in the preclinical and clinical phases of research. A potential therapeutic feasibility of combining such inhibitors with the aim of targeting specific tumor subsets will also be studied further.

## Conclusion

The insilico studies performed using PyRx with Naringin resulted in significant binding affinities against ATM, ATR, CHK1, and WEE1. At the same time, Rutin shows good binding affinity for PARP-1, ATR, and ATM.

SWISS and PkCSM ADMET provide valuable information on the characteristics of Naringin and Rutin. Admet and druglikeness data obtained from insilico are helpful in understanding the lead at an early stage and can be used for further optimization of the molecules in vitro.

The MTT assay showed a range of viability from 75-95% for cells of 2DD and MDA-MB-231 at the concentrations applied. The data indicate that low to medium concentrations (0-160µM) were not toxic.

DPPH assay results indicate a significant level of antioxidant activity at a low concentration (10µM). As a result of MTT and DPPH assays, we selected 10µM concentrations of Naringin and Rutin to test their Free Radical Scavenging activity. Naringin and rutin showed reduced free radical scavenging activity against OH, ONOO, O2− and NO. Analyses of free radicals confirm Narigin and rutin’s antioxidant properties.

In a high alkaline comet assay (pH >13), the molecules demonstrate no DNA damage in normal cells. Human Dermal Fibroblast cells (1×10^5^cells/ml) were pre-incubated with molecules (10 µM) for one hour. The molecules were also pre-incubated with breast cancer cells for one hour. It was observed that the molecules caused 60-70% strand and base damage in the cells. Therefore, naringin and rutin can be considered leading candidates in breast cancer research, although animal and human studies are necessary to confirm their effectiveness.

## Statement of Ethics

“An ethics statement was not required for this study type, no human or animal subjects or materials were used.”

## Author Contributions

Ashwini Badhe- Performed insilico and ADMET studies. Written first draft Pravin Badhe- Performed MTT, DPPH, Free radical assays and Comet assay. Statistical analysis was performed by Pravin Badhe. Vivek Nanaware- Performed editing and proofreading of the manuscript.

## Data Availability

All data analysed during this study are included in this article. Further enquiries can be directed to the corresponding author.

